# The Utility of Red Cell Distribution Width as a Parameter for Calculating Indices of Allostatic Load

**DOI:** 10.1101/085373

**Authors:** Ghalib A Bello, Gerard G Dumancas

**Affiliations:** Dept. of Environmental Medicine & Public Health, Icahn School of Medicine at Mount Sinai, 17 E 102nd St, New York, NY 10029, USA; Dept. of Mathematics & Physical Sciences, Louisiana State University - Alexandria, 8100 Hwy 71 S, Alexandria, LA 71302, USA

**Keywords:** Allostatic Load, Biomarkers, Red blood cell distribution width, Aging

## Abstract

**Background:** Allostatic Load is a construct used to quantify the cumulative burden of exposure to stressors that, over the course of an individual’s life, exert a toll on the body’s physiological functions, increasing risks of various chronic ailments and conditions. Studies attempting to quantify allostatic load have used a variety of clinical biomarkers representing primary and secondary mediators. In this study, we demonstrate the value of including red blood cell distribution width (RDW) among the panel of clinical parameters used to calculate allostatic load.

**Methods:** We develop a novel formulation of allostatic load using RDW and other standard biomarkers. This index is computed using clinical laboratory data from the NHANES study. The predictive validity of the new index for tertiary outcomes (all-cause mortality and physician-assessed health status) is compared to that of the current formulation using Harrell’s C index, ROC analysis and regression-based goodness-of-fit measures.

**Results:** Inclusion of RDW as an allostatic load biomarker yields a significantly improved index. It demonstrates a superior ability to predict mortality, health status and biological age than the standard formulation currently in use.

**Conclusion:** RDW has shown strong correlations with mortality and a broad spectrum of diseases. A review of the existing literature on allostatic load reveals its underutilization in this area, despite being a standard component of blood count panels. This study is the first to demonstrate its usefulness as a potential allostatic load biomarker.

## Introduction

Normal biological functioning in an organism requires maintenance of multiple physiological parameters within narrow ranges, e.g. body temperature, blood pH, blood salinity. This delicate state, known as physiological equilibrium, is disrupted whenever an organism is exposed to environmental stressors. Allostasis is the process by which the body restores homeostatic equilibrium in response to such stressors. Upon exposure to environmental challenges, the body’s stress response mechanisms are triggered and this induces a cascade of biochemical signals designed to restore physiological parameters back to the levels characteristic of p functioning. While useful and indispensable in the short term, these restorative mechanisms, over time, exert negative effects on overall health. Chronic and repeated exposure to environmental challenges and constant triggering of these stress response mechanisms causes eventual deterioration in response efficacy. Over time, this maladaptive cycle gradually leads to dysregulation in multiple physiological systems. The cumulative physiological burden arising from these mechanisms is termed ‘allostatic load’.

The concept of allostasis provides a concise theoretical framework for characterizing and quantifying the overall wear and tear the body endures, which leads to a range of pathologies and chronic conditions. However there has historically been difficulty in operationalizing this concept for practical purposes [1]. The first attempt to quantify allosatic load was by Seeman et al (1997) in the MacArthur Successful Aging Study [2]. In this study, they defined a simple, count-based index using 10 biomarkers spanning multiple physiological systems: metabolic, cardiovascular, HPA (hypothalamic-pituitary-adrenal axis) and sympathetic nervous system activity. Using these biomarkers, allostatic load was then computed using a simple scoring system. For each biomarker, 1 point was assigned if the biomarker value fell outside of the clinically-defined p ranges (e.g.), and 0 otherwise. These points are then summed across all biomarkers to produce an allostatic load score reflective of the underlying cumulative biological burden [2]. This operationalization of allostatic load was validated by testing its association with mortality, declines in physical & cognitive functioning and incident cardiovascular disease. Since Seeman’s landmark paper, the literature has seen a proliferation of alternative formulations of allostatic load composite scores [3, 4]. Ongoing reviews of the ever-expanding literature on this subject point to enormous variation in the way it is operationalized [5].

Red blood cells typically vary in size and shape, but an unusually high degree of variation is evidence of a pathological condition known as *anisocytosis*. Red blood cell distribution width (RDW) is a hematological parameter that quantifies the extent of anisocytosis - it serves as a metric of the degree of heterogeneity in red blood cell volume [6]. RDW was originally used to detect the presence of anisocytosis for differential diagnosis of anemia, however several recent studies have shown that anisocytosis is present in a wide spectrum of conditions, including some with no ostensible relation to hematological disorders [6]. RDW was found to have remarkably high predictive power for mortality, and for several conditions spanning multiple organ systems, e.g. cardiovascular disease [7-10], cancer [11-14], diabetes, liver disease [15, 16], kidney disease [17, 18], respiratory diseases [19], etc. It has also been shown to be associated with oxidative stress, inflammation, poor nutritional status and shortening of telomere length [20]. While it has shown strong and independent associations with, and clinical prognostic value for an extensive panel of conditions, the underlying biological mechanism(s) driving these associations is still unclear. But there is emerging consensus that RDW may be a reflection of severe biological and physiological imbalance [6]. Whether RDW is a causal factor or merely a manifestation of this imbalance is currently unknown. Despite its putative role as a marker of multisystemic dysregulation, no examples exist, to our knowledge, of allostatic load indices that include this marker. This is surprising, given how readily available the RDW parameter is in clinical settings. It is a routinely reported component of standard blood count panels. We also note that RDW has shown associations with multiple health outcomes and biological measures that have been previously linked with allostatic load, e.g. oxidative stress, inflammation, cardiovascular disease, all-cause mortality, telomere length [20, 21], etc. This suggests an interesting, albeit indirect, link between allostatic load and RDW. Also, most studies revealing assocations of RDW with various biological and health outcomes have established its unique role as an independent predictor of these outcomes, i.e. it retains a statistically significant association even after adjusting for other biomarkers associated with the outcome of interest. These observations support the notion that RDW could play an important and independent role in characterizing allostatic load. Therefore in this study, we develop a new allostatic load index that includes RDW and attempt to determine whether, compared to other allostatic load biomarkers currently in use, RDW contributes additional explanatory or predictive power for allostatic load.

## Methods

### Study Population

All analyses described herein are based on clinical and biological data collected by the CDC on a large cohort of the US population. This cross-sectional study, known as the National Health and Nutrition Examination Survey (NHANES) [22], was designed to assess the health and nutritional status of the US population. We used data from the Third NHANES study (NHANES III) which was conducted between 1988-1994.

### Measurement of RDW

RDW is commonly calculated as the coefficient of variation (CV) in red blood cell volume, i.e. the standard deviation of RBC volume dividied by the mean RBC volume (known as Mean Corpuscular Volume [MCV]). This ratio is expressed as a percentage by multiplying by 100:

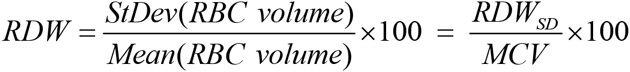

It has recently been shown that the standard deviation of RBC volume (termed RDW-SD) is a stronger measure of mortality than the traditional measure of RDW which divides RDW-SD by MCV [23]. For this reason, we hypothesize that RDW-SD would be a better candidate for inclusion as an allostatic load biomarker and this is the version we use throughout our study.

### Allostatic Load computation

In order to evaluate the potential of RDW-SD as a novel biomarker of allostatic load, we carried out a series of analyses using the NHANES III data described above. We compared a commonly used formulation of the allostatic load score (introduced in Seeman et al. 2008 [24]) with a newer version that contained the same biomarkers but also included RDW-SD. Table 1 summarizes the computation of each allostatic load index. For each biomarker in the allostatic load score, 1 point is assigned if an individual’s value fell outside of the clinically-defined range, and 0 otherwise. These values are then summed to derive a composite score. While multiple alternative allostatic load formulations exist (varying in both biomarker composition and computation method), we chose the formulation developed by Seeman et al. (2008) [24] because it has been used in multiple NHANES studies. As mentioned earlier, in this study we compared Seeman’s original allostatic load index to a new index that iss identical to Seeman’s except for the addition of RDW-SD. To distinguish these 2 formulations, we will henceforth refer to the original formulation as ALI and the new formulation as ALI+RDW.

**Table 1:** Biomarkers in Seeman’s 2008 formulation. (bottom panel shows RDW-SD and the percentile-based cutoff used in this study. For each biomarker, add 1 point if the clinical cutoff is exceeded. Sum points to derive ALI and ALI_+RDW_ scores)

| Variable | High Risk Clinical Cutoff |
| --- | --- |
| Albumin (g/dL) | < 3.8 |
| C-reactive protein (mg/dL) | $\geq 0.3$ |
| Waist-to-hip ratio | > 0.9 (Males)<br>> 0.85 (Females) |
| Total Cholesterol (mg/dL) | $\geq 240$ |
| HDL (mg/dL) | $\geq 40$ |
| A1c (Glycated Hemoglobin [%]) | $\geq 6.4$ |
| Resting Pulse Rate (beats/min) | $\geq 90$ |
| Systolic BP (mmHg) | $\geq 140$ |
| Diastolic BP (mmHg) | $\geq 90$ |
| New Variable | High Risk cutoff (75th percentile) |
| RDW-SD | > 12.2 |

### Assessment of Allostatic Load Indices

To gauge the quality of the new allostatic load formulation relative to that of the commonly used version, we followed the approach of Seplaki et al. (2005)[25] - comparing their predictive ability and explanatory power for tertiary outcomes of increased allostatic load. We use mortality and physician-assessed health status as outcome measures. Both of these outcomes are strong reflections of allostatic load and underlying physiological burden. Mortality is the ultimate outcome of the cumulative physiological dysregulation that arises from elevated allostatic load. The first operationalization of allostatic load developed by Seeman et al. (1997) [2] used mortality was one of the outcomes for assessing the predictive validity of the newly formulated construct. Multiple studies since then have used all-cause mortality as a surrogate metric for judging new formulations of allostatic load. In this study, mortality data was obtained from the NHANES III National Death Index (NDI) linkage files. These files provide information about the all-cause mortality status and survival times (up to December 31, 2011) of NHANES III participants [22]. The predictive power of ALI+_RDW_ and ALI for all-cause mortality was compared using Harrell’s concordance index (Harrell’s C for short) [26]. Harrell’s C is an U-statistic that estimates the probability that, for a randomly selected pair of individuals, the one with the higher allostatic load score will have shorter survival time, and vice versa. A better mortality predictor should therefore have higher Harrell’s C. In addition, ROC (Receiver Operating Characteristic) analysis for 10-year mortality was performed by dichotomizing the continuous survival times into binary categories, i.e. the cohort was divided into those who died within 10 years from baseline (the year of NHANES participation) and those that survived past the 10-year mark. The AUC measures for ALI and ALI+rd_W_ were compared (using DeLong’s test) to determine which had better predictive value for long-term mortality.

Besides mortality, ALI and ALI+rdw were also compared on their ability to predict physician-assessed health status. A subset of NHANES participants underwent standardized, comprehensive physical examinations performed by board-eligible physicians. These physicians then assigned health status rankings to each individual based on their overall impression of their health. A simple rating scale was used, with health rankings of Excellent, Very Good, Good, Fair and Poor. For analytical purposes, we dichotomized this health status score by categorizing individuals into two groups: 1) Excellent, Very Good, or Good health; 2) Fair or Poor health. ROC analysis was used to compare AUC measures of ALI and ALI+_rdw_ with respect to their ability to predict the binary health status.

Studies have established that aging is inextricably linked with allostatic load [27]. Allostatic load accumulates as individuals age, and it has been proposed as a model of how lifetime exposure to environmental stressors leads to the gradual declines in physiological function that accompany aging. Indeed, there is a growing corpus of evidence linking allostatic load to a number of age-related conditions like frailty [28], dementia [29] and diminished physical functioning [30]. It is noteworthy that measures of Biological Age [31] are constructed using many of the same biomarkers used in allostatic load indices. We therefore reasoned that age is a strong proxy for allostatic load, and since the latter increases with age, a good measure of this construct should show a strong associational gradient with age. Using multivariate linear regression models, ALI and ALI+_rdw_ were compared with respect to their explanatory power for chronological age. Separate models were fitted for ALI and ALI+_rdw_. In each model, age was used as the dependent variable while each index was included as an explanatory variable. Both models were adjusted for sex and race/ethnicity. R^2^ and AIC (Akaike Information Criterion) were compared for the 2 models in order to determine which provided a better fit for age.

### Expanded set of allostatic load biomarkers

Since the first operationalization of allostatic load was proposed by Seeman and costring-names in 1997 [2], several alternative formulations have been developed and utilized. In this study, we focused only on the version Seeman et al. (2008) [24] developed for use with NHANES data [24]. While this formulation has been heavily utilized in the literature, the 9 biomarkers it is comprised of represent only a subset of the biomarkers that have been used across the cumulative body of work on allostatic load. Thus we were interested in assessing the performance of RDW-SD in the presence of a more comprehensive panel of allostatic load biomarkers. The goal of this assessment is to determine whether RDW-SD is a truly novel allostatic load biomarker, in the sense that it provides information about physiological dysregulation that no *currently used* allostatic load biomarker does. In other words, we tried to address the question: are there currently or previously used biomarkers of allostatic load that provide the same information that RDW-SD does, thereby rendering it redundant? To answer this question, we reviewed the literature on allostatic load studies utilizing NHANES data. We compiled a list of 22 biomarkers that have been used in various allostatic load formulations across multiple studies (see table 1). For each one of the outcome variables (mortality, health status and age), we separately fitted multivariate models that included the entire pool of 22 biomarkers and RDW-SD as explanatory variables, adjusting for demographics. Each of these models had mortality, health status or age as dependent/outcome variables. For the mortality model, we used a Cox Proportional Hazards Regression. For health status, we used both ordinal regression (for the raw health status score) and logistic regression models (for the dichotomized version of the raw score). And for age, we used multivariate linear regression. In all these models, we assessed the statistical significance of RDW-SD in the presense of the other 22 allostatic load biomarkers to determine if RDW-SD exhibits independent associations with each of the outcome variables. If so, this would imply that RDW-SD captures aspects of allostatic load that other allostatic load biomarkers do not.

All analyses were carried out in SAS (Cary, NC) [32] and R [33].

**Table 2:**
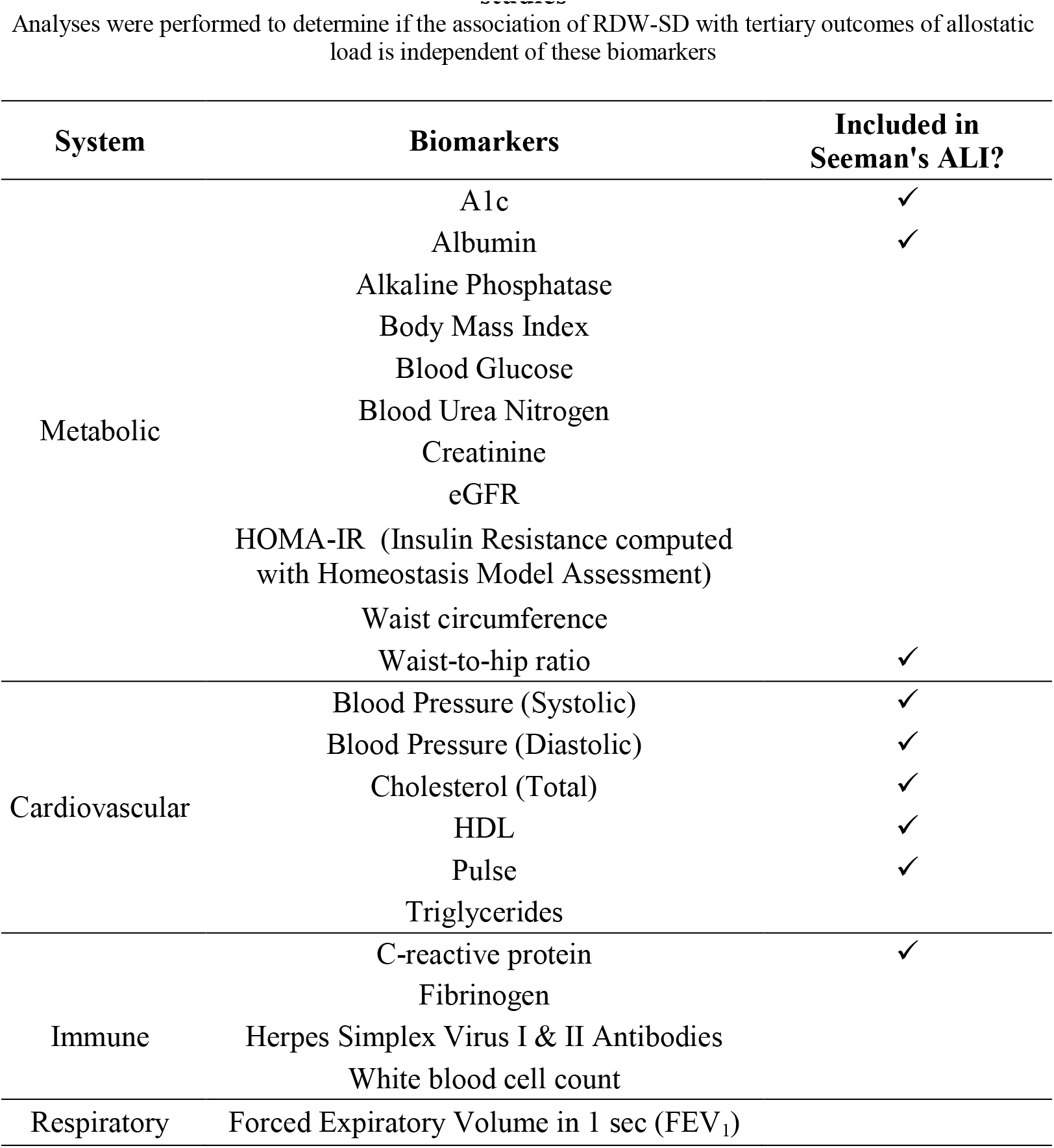
Expanded set of biomarkers that have been used in various NHANES studies. Analyses were performed to determine if the association of RDW-SD with tertiary outcomes of allostatic load is independent of these biomarkers

## Results

Our analytical sample was restricted to individuals over the age of 18 who had complete (nonmissing) data on demographics (age, sex, race), all 9 biomarkers comprising the ALI, RDW, and health outcomes used in this study (mortality and physician-rated health status). A total of n=14,578 NHANES III subjects remained after the above selection criteria were imposed. Table 3 summarizes descriptive statistics on various measures for this sample. The age range was 18-90 years, with a median of 42 years. Nearly 53% were female.

**Table 3:**
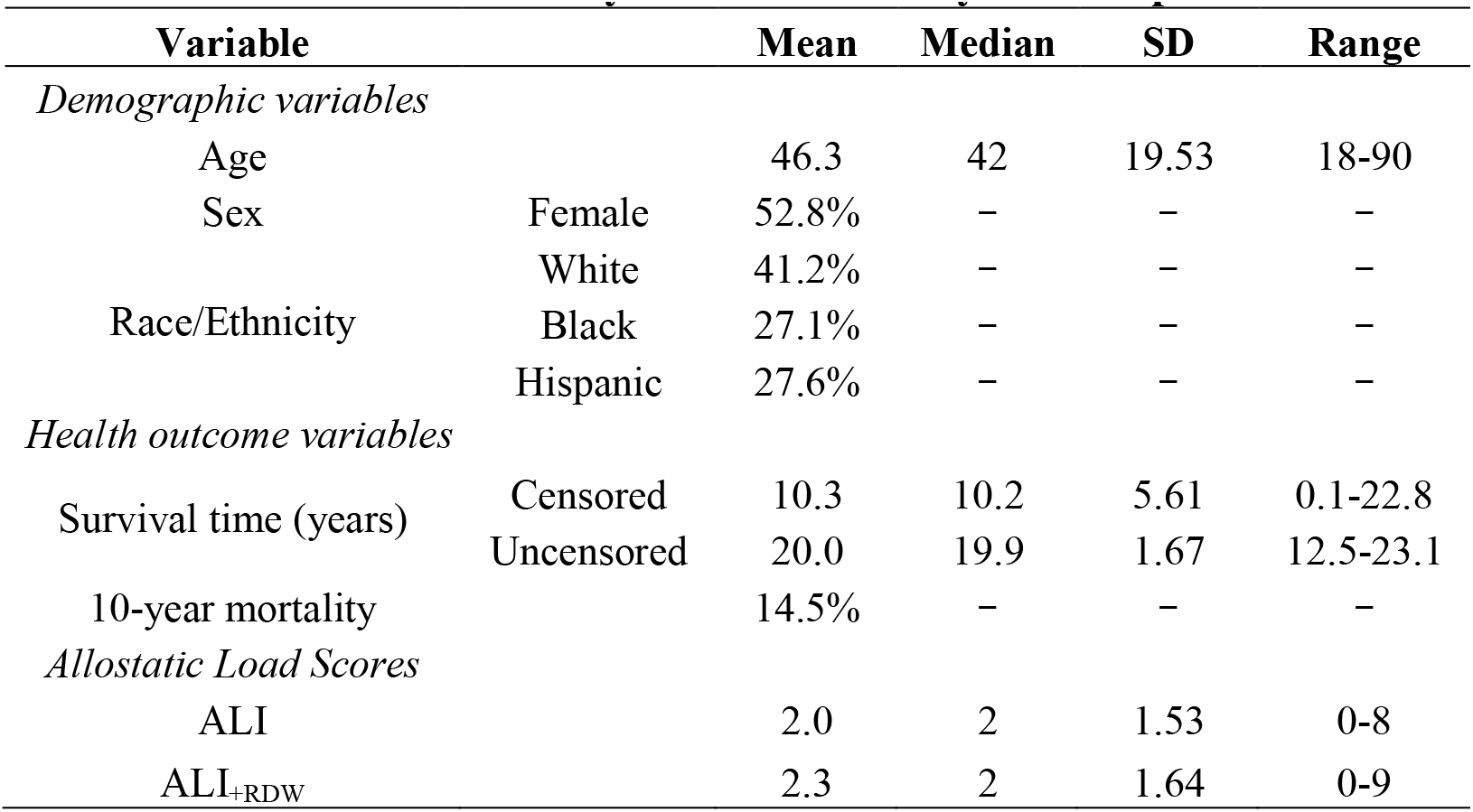
Summary statistics on analytical sample.

### Allostatic load scores

Allostatic load scores were calculated as outlined in Table 1. The biomarkers from the original ALI formulation used clinical cutoffs, while the RDW-SD used a quartile-based cutoff (values exceeding the 75th percentile for adults are considered high). We resorted to using a quartile-based cutoff since RDW-SD does not have any universally accepted clinical cutoffs [23]. Descriptive statistics on allostatic load scores in the study sample are summarized in Table 1. In the analytical sample, the mean ALI score was 2, while the mean ALI+_rdw_ score was 2.3. For both scores, the median value was 2.

### Mortality and Allsotatic load scores

In the analytical sample, the median survival time overall (censored or uncensored) was (Table 1). Mortality within 10 years of NHANES participation was observed in 14.5% of the sample. For the ALI, the Harrell’s C index for continuous survival was 0.7 (95% CI: [0.686-0.715]), compared to.73 for ALI (95% CI: [0.713-0.741]). A similar trend was also observed with 10- year mortality, with ALI+rdw yielding a higher AUC (0.75 [95% CI: (0.740-0.761)]) than ALI (0.72 [95% CI: (0.705-0.727)]). This difference in AUC was statistically significant (p <.0001).

For the dichotomized health outcome, the AUC for ALI_+RDW_ (0.79 [95% CI: (0.777-0.790)]) was also statistically significantly higher (p <.0001) than that of ALI (0.76 [95% CI: (0.750-0.774)]). The significant increase in predictive power for both mortality and health status are remarkable given that they are due to the addition of just one biomarker to the original allostatic load score. Interestingly, we found that RDW-SD by itself predicts 10-year mortality with an AUC of 0.71, statistically equivalent to the AUC for the ALI. Thus, we see that just on its own, RDW-SD performs as well in predicting mortality as the ALI composite, which comprises accumulated information from 9 diverse biomarkers.

### Association with Age

Table 4 below summarizes the adjusted R^2^ and AIC for sex-and race-adjusted models of age using either ALI or ALI+rd_W_. Both ALI and ALI+rd_W_ exhibited significant associations with age in their respective models (p<.0001 in both cases). However, ALI+rd_W_ showed superior performance based on both goodness-of-fit measures, implying its greater explanatory power for age than ALI.

**Table 4:** Goodness-of-fit measures.

|  | **Adjusted $R^2$** | **AIC** |
| --- | --- | --- |
| ALI | 0.295 | 122895.8 |
| ALI<sub>+RDW</sub> | 0.333 | 122088.5 |

### Expanded biomarker pool

Due to the inclusion of the expanded set of 22 biomarkers, the original analytic sample size (n=14,578) was reduced to n=4098. This reduction was due in large part to the fact that a couple of the biomarkers in the expanded pool (particularly fibrinogen and Herpes I & II antibodies) were only measured for a fraction of NHANES III examinees. In the models for mortality (continuous survival & 10-year mortality), health status and age, RDW-SD maintained a significant p-value (p<.0001) in the presence of the entire pool of 22 biomarkers, indicating its independent association with these outcomes.

## Discussion

Seeman and co-workers in their groundbreaking 1997 study [2] developed the first operationalization of allostatic load with 10 biomarkers. In a more recent 2008 study using NHANES data (Seeman et al. 2008) [24], they developed a different allostatic load formulation due to the lack of availability of certain biomarkers in the NHANES data. This formulation was subsequently adopted by other researchers [34-36]. In this study, we have shown that this formulation can be significantly improved upon by the inclusion of RDW-SD. The added predictive ability of RDW-SD for tertiary outcomes of physiological dysregulation (mortality, health status) indicates its potential as a new allostatic load biomarkers. Further, its independent, significant association with chronological age shows that it could be a promising candidate as a biomarker of biological aging. Our study corroborates the growing evidence regarding the strong relationship between age and RDW.

Beyond providing what is arguably a better allostatic load score than the currently used formulations, this study is the first to highlight the possible link between anisocytosis and allostatic load. Increased RDW has been shown to be related to many conditions that allostatic load has been linked to. We propose that a common link between the two may be inflammation, and hope that this study serves as a catalyst for further investigations into this connection. Currently, most allostatic load formulations include a few biomarkers of inflammation (primarily C-reactive protein [CRP]). We propose inclusion of RDW-SD as another inflammatory allostatic load biomarker. Interestingly, RDW and CRP share many similarities, and studies have shown the two are significantly correlated [37]. Our study results prove that RDW still retains predictive ability even in the presence of CRP, demonstrating that despite its correlation with CRP, it provides additional, non-redundant explanatory power for mortality, health status and age. In addition, RDW is in many cases more readily available than CRP, since it is a component of standard blood count panels. CRP is more expensive to measure and does not belong to any standard clinical diagnostic test panels. For calculating allostatic load, we propose including RDW in lieu of CRP in settings whereby measurements of the latter are unavailable.

Our study has a few limitations. Firstly, we compared the allostatic load formulations using only mortality, health status and age. However, there are other tertiary outcomes that could be used, e.g. frailty, cognitive decline, physical function decline, cardiovascular disease incidence, etc. Future studies will focus on using a wider set of outcomes. Secondly, our analysis is restricted to NHANES data, which historically has not collected a few key biomarkers used for operationalizing allostatic load, particularly primary mediators such as nervous system biomarkers (cortisol, catecholamines and dihydroepiandrosterone sulphate (DHEA-S)). These biomarkers play a central role as mediators of downstream effects of the primary stress response [38]. In addition, there are other biomarkers that have been used in other studies which NHANES data limitations prevented us from including in our analyses. So the 22 biomarkers examined in this study by no means constitute an exhaustive list of biomarkers that have been used in various formulations of allostatic load. We do, however, note that the validity of some of these biomarkers as allostatic load components has been questioned [39]. Domain experts have warned that, increasingly, many of the alternative allostatic load formulations being proposed sometimes include biomarkers that have little to do with the concept of allostatic load. Goldman [39], for example, claims some new measures are largely ‘atheoretical’, with too much focus concentrated on the performance of biomarkers (ability to predict various outcomes) as opposed to their overall relevance to the notion of allostasis. We agree that the appropriateness of a novel biomarker as a potential component of allostatic load should be judged based on both predictive/explanatory power and theoretical relevance to the concept of allostasis. In this paper, we have presented a case to demonstrate that RDW satisfies these two criteria of predictive validity and concept validity. We have shown that RDW provides supplemental information about physiological dysregulation (beyond what many currently used allostatic load biomarkers provide), and that it appears to be a marker of physiological processes relevant to the conceptualization of allostatic load.

